# Democratizing Agentic Access to Bioinformatics and Biopharmaceutical Databases and Analyses with BioMCP-TS

**DOI:** 10.64898/2026.09.03.749120

**Authors:** Ye Yuan

## Abstract

Practitioners want to give AI agents direct access to biomedical databases and analyses, but today that means choosing between dependency-heavy skill packages, data-lookup-focused server tools, and closed vendor platforms. BioMCP-TS removes the choice: one open-source server that installs with a single pinned command and verifies its own setup. An agent working through BioMCP-TS can search and cross-reference 50+ bioinformatics, pharmaceutical, and patent databases (genes, variants, drugs, diseases, literature, clinical trials, patents, functional genomics, and structures) and, uniquely among open bioinformatics servers, run heavyweight analyses in-process as WebAssembly: Bioconductor differential expression and htslib genomics operations with no R installation, C toolchain, or containers on the host, plus read-only SQL over curated local databases. We demonstrate these strengths with seven practical cases spanning drug-target due diligence, translational intelligence, GWAS follow-up, dependency analysis, cohort genomics, and differential expression, ranging from a first federated lookup through target-disease landscapes to published-structure shortlists and reproducible RNA-seq results, each answered end-to-end within minutes. The server, recorded transcripts, and the full problem set are available at https://github.com/yeyuan98/biomcp-ts (npm: `biomcp`).

## Introduction

Biomedical knowledge is spread across dozens of public APIs. Gene annotations, variant interpretations, drug targets, disease associations, literature, clinical trials, expression resources, and sequence archives each expose their own endpoints, identifiers, and rate limits. Whole careers are spent bridging them with scripted pipelines, workflow systems such as Galaxy (Afgan et al., 2018), and aggregator APIs such as the biothings services behind MyGene and MyVariant (Wu et al., 2013; Xin et al., 2015; Lelong et al., 2022). Practitioners increasingly want their AI agents to do this bridging on demand, in natural language, without a data engineer in the loop.

The Model Context Protocol (MCP) has quickly become the standard way to expose such tools (Anthropic, 2024). Under MCP, a host application (a coding assistant, an analysis agent, or an agent CLI) launches a small server program locally and calls the functions, called tools, that the server advertises; each tool ships with a machinereadable schema of its arguments, so the agent can discover and compose capabilities without bespoke wrappers, and the conversation runs over local standard input and output (stdio). Life-science MCP deployments and perspectives are accumulating (Flotho et al., 2026; Widjaja et al., 2025; Saez-Rodriguez, 2025).

Equipping an agent for biomedical work currently means choosing among three integration styles, each with complementary weaknesses (Table 1). All-in-one vendor science platforms such as Claude Science and the Rosalind Workbench (Anthropic, 2026; OpenAI, 2026) offer polished, tightly integrated scientific workbenches, but they are closed-source and bound to one vendor’s models, data partnerships, and release cycle. Skill- and plugin-based approaches wrap local commandline programs and library workflows, so they can reach the full analysis stack, including Bioconductor (Love et al., 2014; Robinson et al., 2010; Ritchie et al., 2015) and the C-based htslib suite (Li et al., 2009; Li, 2011; Quinlan and Hall, 2010); that flexibility is paid for in installation and maintenance, since each host needs matching Python or R environments, and skills are less machine-discoverable than schema-advertised tools (Anthropic, 2025; jaechang-hits, 2026; GPTomics, 2026). MCP-based bioinformatics servers (Widjaja et al., 2025; Genomoncology, 2025) install in seconds and are lightweight and self-describing, but the ones available today concentrate on API federation and data lookup; to our knowledge, none ships in-process Bioconductor or htslib compute. WebAssembly builds of the core tooling exist and are proven in browsers (biowasm project, 2019; Stagg and Henry, 2023; Ji et al., 2024), but this compute path has not been wired into a bioinformatics agent server.

**Table 1.** Integration styles for giving AI agents biomedical data and tools. Rows are qualitative summaries of the cited representatives’ documented scopes; competitor attributes were not independently benchmarked. A tool is a function the agent can call; a host is the agent application.

|  | Vendor science platforms | Skill-/plugin-based agents | MCP-based bio servers | BioMCP-TS |
| --- | --- | --- | --- | --- |
| Representatives | Claude Science (Anthropic)<br>Rosalind Workbench (OpenAI) | Agent skills (Anthropic)<br>SciAgent-Skills (jaechang-hits)<br>bioSkills (GPTomics) | BioinfoMCP<br>BioMCP (genomoncology) | this work |
| Openness | closed-source, vendor-bound | open, community libraries | open, single servers | open (Apache-2.0) |
| Host installation | none (hosted) | Python or R environments per skill | one server process | one server process |
| Data federation | vendor-curated | case by case | moderate (per-domain) | 50+ databases, 41 core tools |
| Analysis compute | vendor-side, not inspectable | host-side, dependency-heavy | not provided | in-process WebAssembly |

BioMCP-TS (https://github.com/yeyuan98/biomcp-ts; npm package `biomcp`, where npm is the Node.js package registry) closes that gap, and the point is democratization: one install command, zero configuration, and any practitioner’s agent can query 50+ upstream databases through 41 core tools, then optionally load plugin sets for SQL analytics and for analyses running as WebAssembly inside the server process, with no R installation, C toolchain, or container runtime on the host.

This report showcases the server through its use. Seven recorded practical cases span the application lanes. An onboarding case goes from a clean machine to a first federated answer (Case 0). Three exercise the federation: an SBDD target due-diligence cross-reference for IL-23R (Case 1), a disease-entry translational-landscape map for sickle-cell gene therapy (Case 2), and a GWAS-hit functional-genomics triage at SORT1 (Case 3). Three exercise the optional plugins: DepMap CRISPR dependency analysis over SQL (Case 4), cohort-scale genomics compute chaining BAM, BED, and VCF tools on real human sequencing fixtures (Case 5), and differential expression of a public 27,179-gene mouse dataset (Case 6). Every case was executed against pinned release v1.1.0 with the problem, the exact prompt, the tool-call sequence, and the answer recorded verbatim; the full problem set ships as a published dataset for reproduction. The next section presents the cases; the System design section then explains how the server delivers them and how it is tested; limitations close the report. Table 1 positions BioMCP-TS against the integration styles a practitioner can choose today.

## 2. Practical cases

Every case below was recorded live against pinned v1.1.0: a fresh server process, a natural-language prompt, and the full tool-call sequence, answer, and transcript preserved verbatim in the published problem set (Availability). The cases are chosen to showcase what an agent can *get done* through BioMCP-TS; the answers and the identifiers below are the recorded outputs, not illustrations.

### 2.1 Case 0: Install, verify, and answer (onboarding)

A practitioner with only Node.js wants a working biomedical server for their agent. The README’s pinned install command sets the server up on a clean machine, its built-in self-check confirms the installation is healthy, and the agent’s first federated lookup already returns a usable answer: SORT1 (sortilin 1), GRCh38 chr1:109,309,568–109,397,967, a VPS10-family trafficking receptor, coordinates that recur in Case 3’s deep dive (Fig. 1). The whole journey, from empty machine to first answer, is a single short session.

**Figure 1.**
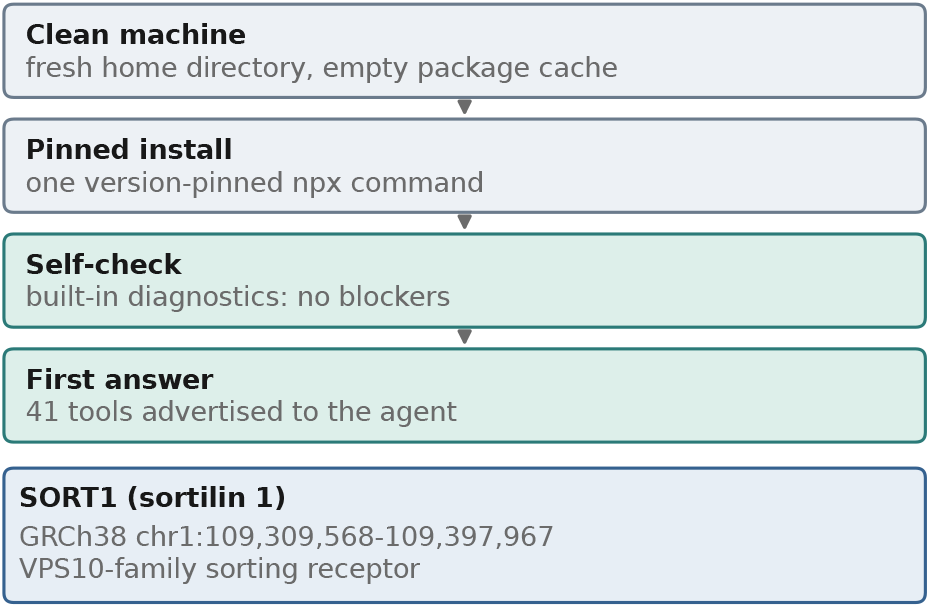
Case 0 (onboarding): clean machine → pinned install → self-check → first federated answer, recorded at v1.1.0.

### 2.2 Case 1: SBDD target due diligence for IL-23R (core tools)

*Prompt*. “I want to start an SBDD program against the interleukin-23 receptor (IL-23R). Give me the target’s biological mechanism, existing drugs targeting the IL-23 axis with FDA status and trial status, the key literature, relevant patents, and all published IL-23R structures with PDB IDs.”

A medicinal chemist’s pre-docking cross-reference, normally assembled across six portals, arrives in one session (Fig. 2). The answer separates what reads like success from what actually is: no approved drug targets the receptor itself; the approved IL-23-axis biologics (risankizumab, guselkumab) block the ligand, while receptor-directed agents have stalled (brazikumab, terminated) or are still recruiting (eltrekibart). The key structural paper is surfaced (PMID 29287995, Immunity) and the patent estate mapped (the Protagonist binding-protein family and related estates). One lesson was recorded verbatim: PDB text search returns false positives, while the identifier path recovers the real docking-ready structures, 5MZV (2.8 Å), 6WDQ (3.4 Å), and 8OE4 (cryo-EM).

**Figure 2.**
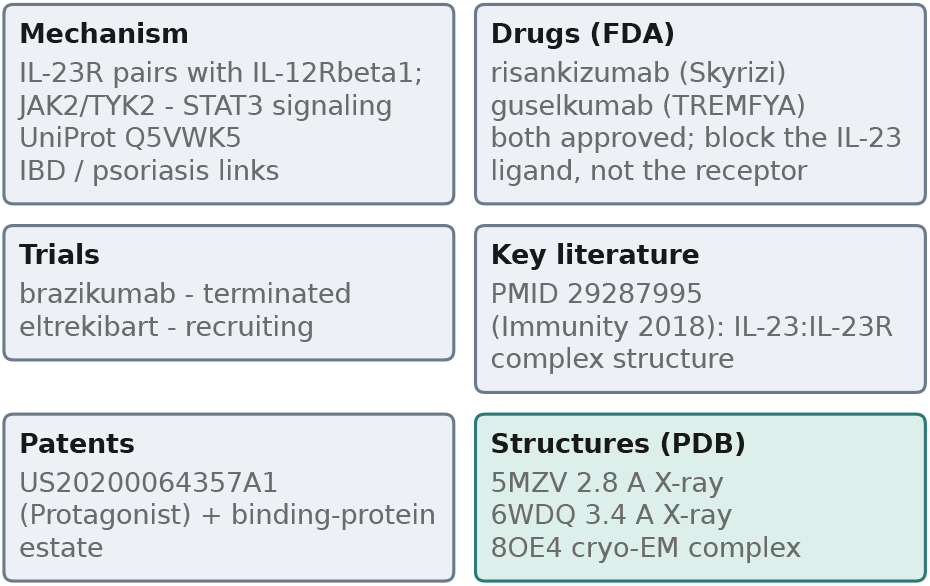
Case 1 (SBDD due diligence, IL-23R): the recorded answer as a card per evidence class (mechanism, FDA drugs, trials, key literature, patents, and the docking-ready structures recovered via the identifier path after the PDB text-search miss).

### 2.3 Case 2: Translational landscape of sickle-cell gene therapy (core tools)

*Prompt*. “Map the translational landscape of sickle-cell gene therapy: the current condition-level trial picture, recent key literature, and the patent estate including foundational prior art.”

Four calls produce a complete landscape for a business-development or translational read (Fig. 3): the ontology anchor (MONDO:0011382), the condition-level trial picture with its supportive-care mix flagged explicitly, five deduplicated recent articles across three literature backends, and a patent estate of 11,415 US document families whose co-citation mining surfaces the field’s foundational prior art, WO1995/011699 (the 1995 PCT behind HbF-induction therapy), exactly the anchor a freedom-to-operate scan needs.

**Figure 3.**
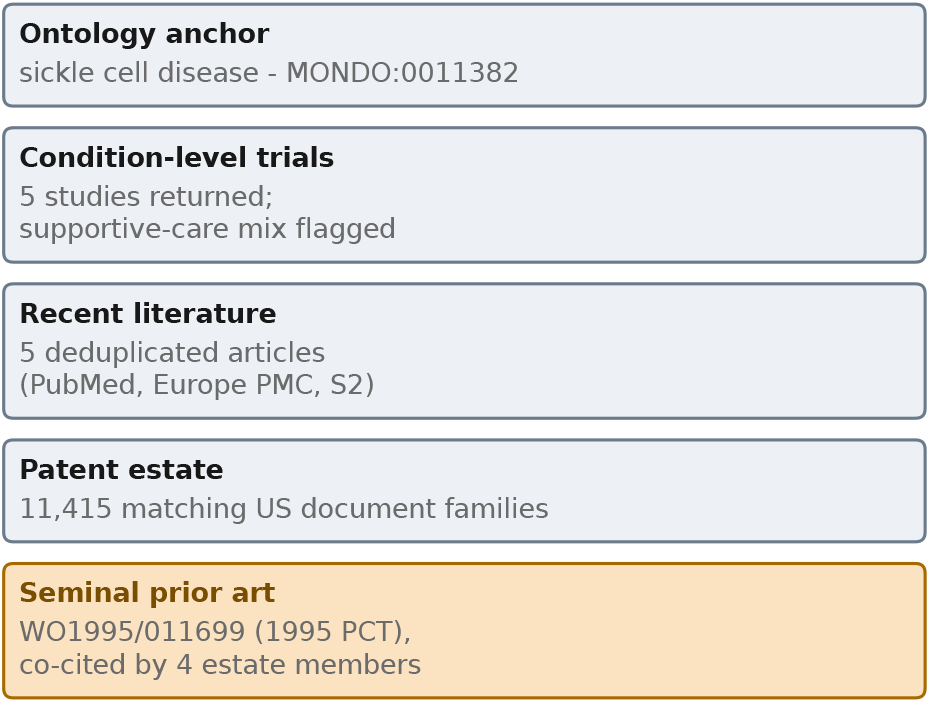
Case 2 (sickle-cell gene therapy): the recorded landscape (ontology anchor, condition-level trials, deduplicated literature, the patent estate, and the co-citation-mined seminal prior art WO1995/011699).

### 2.4 Case 3: Functional-genomics triage of a GWAS hit (core tools)

*Prompt*. “I have a GWAS hit at 1p13.3 linked to LDL cholesterol; the index variant is rs12740374 near SORT1. Before planning wet-lab follow-up, characterize the functional genomics landscape: confirm SORT1 coordinates and transcripts, its tissue expression and liver eQTL signal, its mouse ortholog, available GEO/SRA datasets for the SORT1 enhancer, and the reference sequence around the lead variant.”

The recorded answer tells a wet-lab scientist whether the locus is worth committing to (Fig. 4): the strongest liver cis-eQTL sits at the index variant itself (*p* ≈ 10^−65^) with no coding consequence, a regulatory mechanism rather than a protein change; liver expression is low while artery and brain dominate; the mouse ortholog is one-to-one (91.2% identity); public ENCODE MPRA datasets and ATAC runs provide ready assay systems; and a 201-bp reference slice centered on the lead variant is a ready primer/reporter substrate. SORT1 comes out a credible, testable regulatory target.

**Figure 4.**
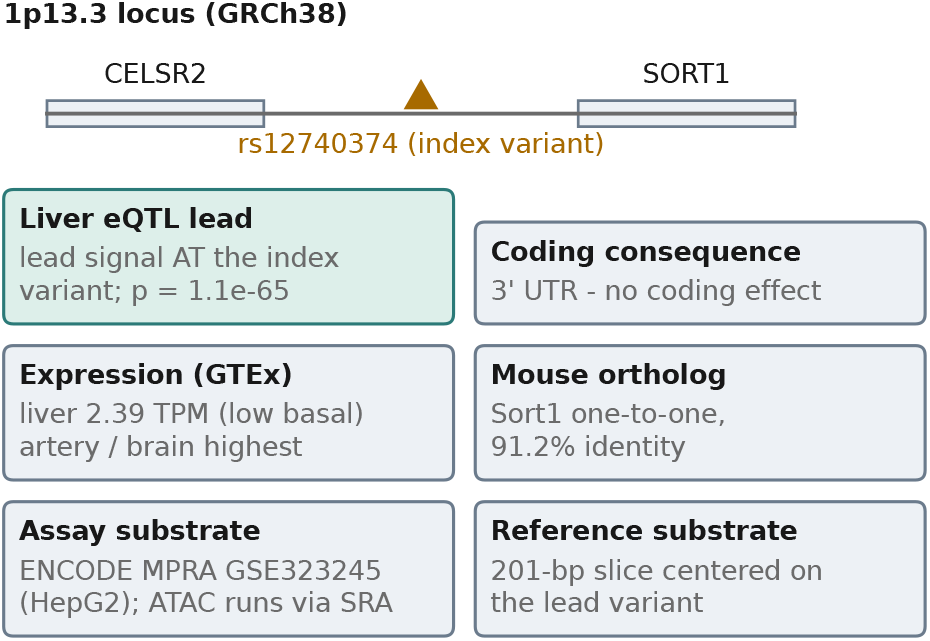
Case 3 (SORT1 / rs12740374): the 1p13.3 locus with the index variant between CELSR2 and SORT1, and the recorded evidence for a regulatory, testable target (liver eQTL lead at the index variant, no coding effect, GTEx expression profile, mouse ortholog, assay datasets, and the reference substrate).

### 2.5 Case 4: DepMap CRISPR dependency analysis over SQL (database plugin)

*Prompt*. “Which 10 DepMap cell lines are most dependent on KRAS, and is that dependency shared across lineages? Summarize from the staged DepMap 26Q1 database.”

With the nine pinned DepMap 26Q1 files staged once (CAPTCHA-gated portal; CC BY 4.0 (Broad Institute DepMap Portal, 2026)), the ETL pipeline builds a 5.62 GB SQLite reference database in a couple of minutes, and two parameterized SQL queries return the answer immediately (Fig. 5): the dependency is dominated by pancreatic, biliary, and bowel lines (AsPC-1 at effect −4.46) but spreads across twelve lineages, a long tail that a notebook analysis would likely miss.

**Figure 5.**
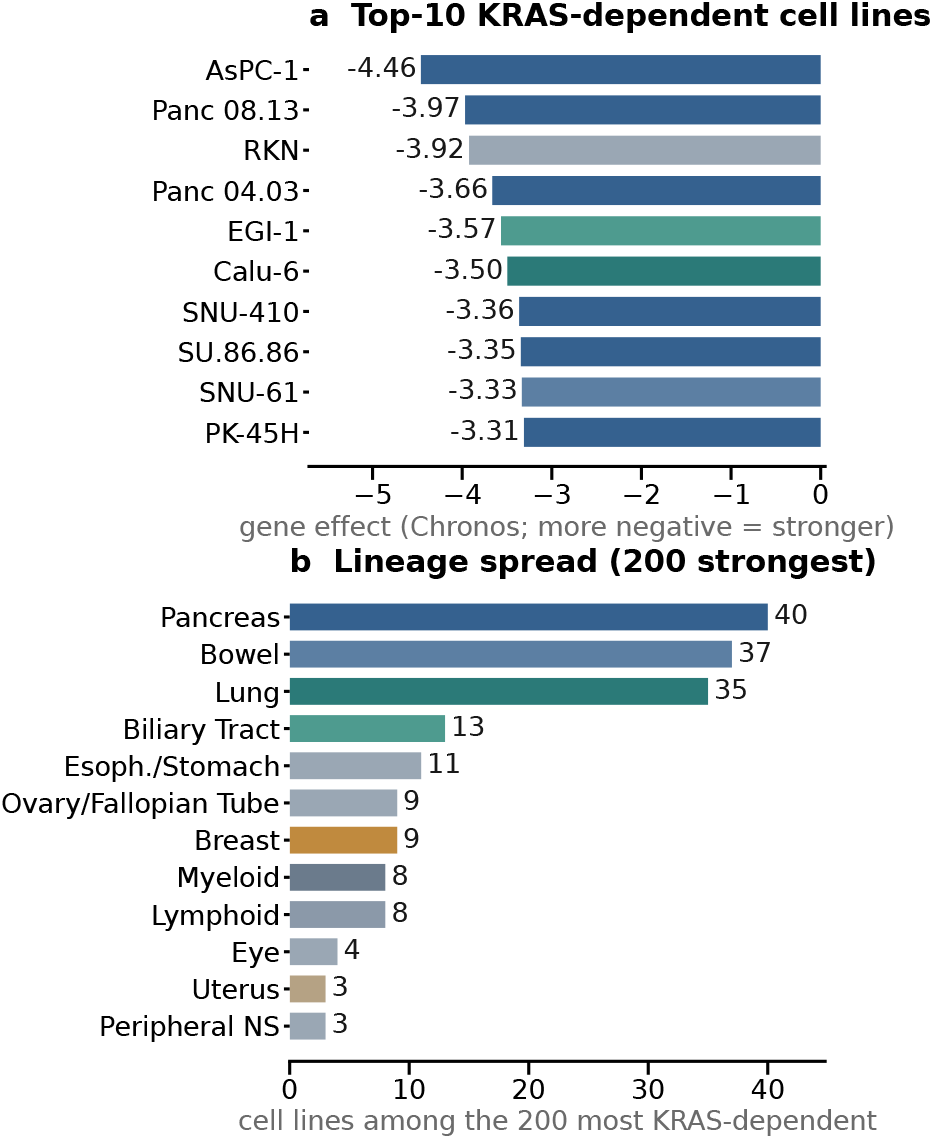
Case 4 (DepMap 26Q1 over SQL), recorded at v1.1.0: (a) the ten most KRAS-dependent cell lines (Chronos gene effect; more negative = stronger), colored by Oncotree lineage; (b) lineage distribution of the 200 strongest dependencies.

### 2.6 Case 5: Cohort triage on real human sequencing data (biowasm plugin)

*Prompt*. “Assess NA12878 coverage over the chr20 JAG1 target from the BAM, derive confidently-covered intervals (≥8*×*) with BED merge/intersect against the panel BED, then extract rare (AF*<*0.01) cohort variants from the 2,504-sample chr22 VCF in CHEK2 and rank the 10 rarest as a final variant list.”

A genomics analyst chains BAM, BED, and VCF operations on half a gigabyte of real human sequencing fixtures with no htslib on the host, in one short session (Fig. 6). The coverage verdict and interval funnel (96 raw → 93 merged → 26 panel intervals) come out with per-base honesty: this NA12878 fixture is low-coverage, and only 12% of the panel target reaches 8*×*, stated rather than tuned away. The CHEK2 window of the 2,504-sample cohort VCF yields 3,211 rare variants (AF*<*0.01), and the final list ranks the ten rarest, all of them unobserved alternate alleles in this cohort. When a request would have streamed the entire cohort VCF (an operation on the order of ten minutes), the server’s large-input gate declines it and suggests slicing instead, the documented workaround the session then takes.

**Figure 6.**
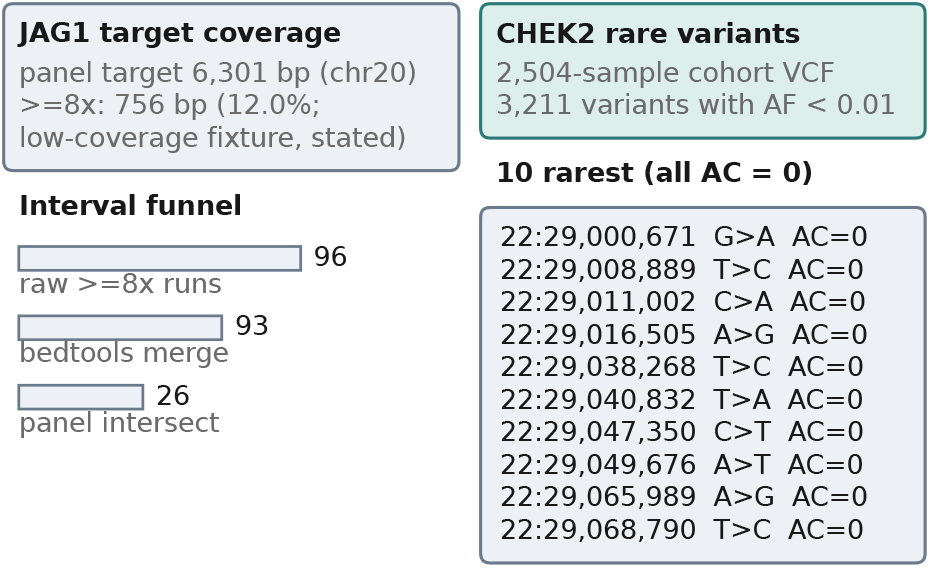
Case 5 (cohort triage): the recorded JAG1 coverage verdict and interval funnel, and the CHEK2 rare-variant extraction with the ten rarest cohort variants (all allele count zero in 5,008 chromosomes).

### 2.7 Case 6: Differential expression of a public mouse dataset (webR plugin)

*Prompt*. “Download the GSE60450 lactation counts, run DESeq2 with design `∼cell + status` contrasting lactation vs virgin, report the top mouse genes and pathway enrichment of their human orthologs.”

A standard Bioconductor workflow (Love et al., 2014) runs endto-end with no R installation anywhere: DESeq2 executes inside the server on the full 27,179 *×* 12 count matrix, the top up-regulated genes are exactly the milk-protein cluster biology expects (Csn1s2b, Wap, Glycam1, Csn2; Fig. 7a), and, because the prompt says so, the cross-species step happens in the same session: fifteen orthology lookups map the top mouse genes to their human orthologs, with the casein-cluster genes correctly reported as having no one-to-one human ortholog (Fig. 7b). Pathway enrichment of the mapped orthologs lands on ERBB4 signaling and connexin oligomerization, a lactation-program signal (Fig. 7c). (An earlier unqualified “report pathway enrichment” prompt derailed this step against human-locked gene tools; the refined prompt and both recorded transcripts are published in the problem set.)

**Figure 7.**
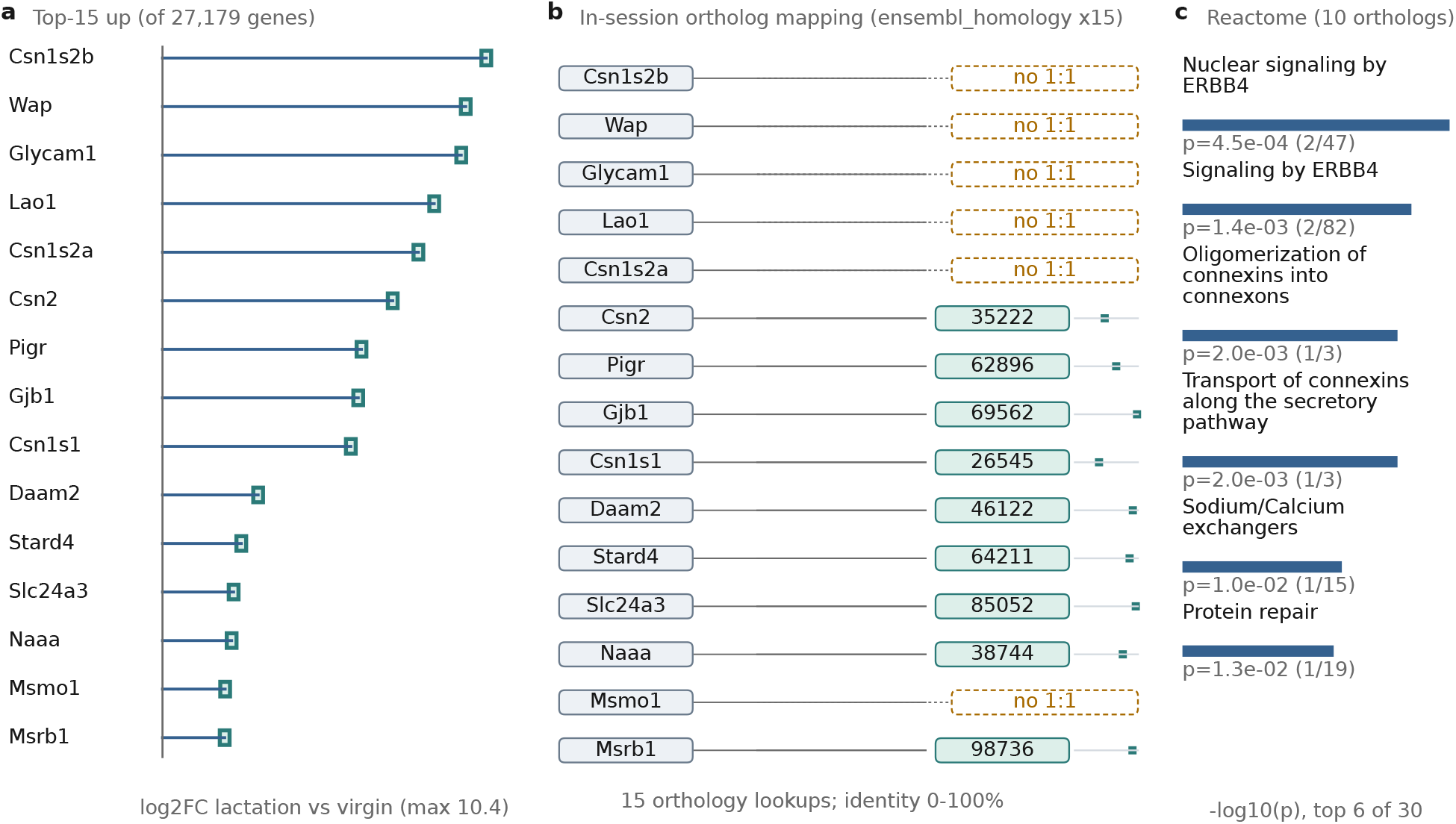
Case 6 (GSE60450 differential expression), recorded at v1.1.0. (a) Top-15 up-regulated genes (the symbol-probed set of the 27,179-gene matrix), lollipop length = log2 fold-change lactation vs virgin (every gene in the set has adjusted *p* ≤ 1 *×* 10^−24^). (b) The in-session cross-species step: 15 mouse genes mapped through orthology lookups to 10 human orthologs (Ensembl gene IDs shown as their unique suffixes; dot on the mini-axis = protein identity 0–100%), while 5 milk-cluster genes terminate at the explicit “no 1:1” state: true database zeros, reported as observed. (c) Reactome over-representation of the 10 orthologs (top 6 of 30 returned terms).

## 3. System design and implementation

This section explains how the server delivers the cases above, and how it is tested; it is deliberately separated from the practical cases so a practitioner can read the cases first and dive into design only when needed.

### 3.1 Overview

All claims refer to BioMCP-TS v1.1.0 (npm biomcp@1.1.0, tarball SHA-1 `fbb086fa`, git tag v1.1.0 equals the PR-merge commit `24c174f`): a TypeScript server (about 26,400 non-test lines; 53,400 including tests) on Node.js ≥ 22.13 (OpenJS Foundation, 2026) and the MCP TypeScript SDK 1.29, exposing a stdio transport with zero configuration and a single runtime dependency (`undici`). Tool arguments are validated against JSON schemas advertised through the protocol. Figure 8 shows the architecture; Figure 9 shows the full tool inventory parsed directly from the registry.

**Figure 8.**
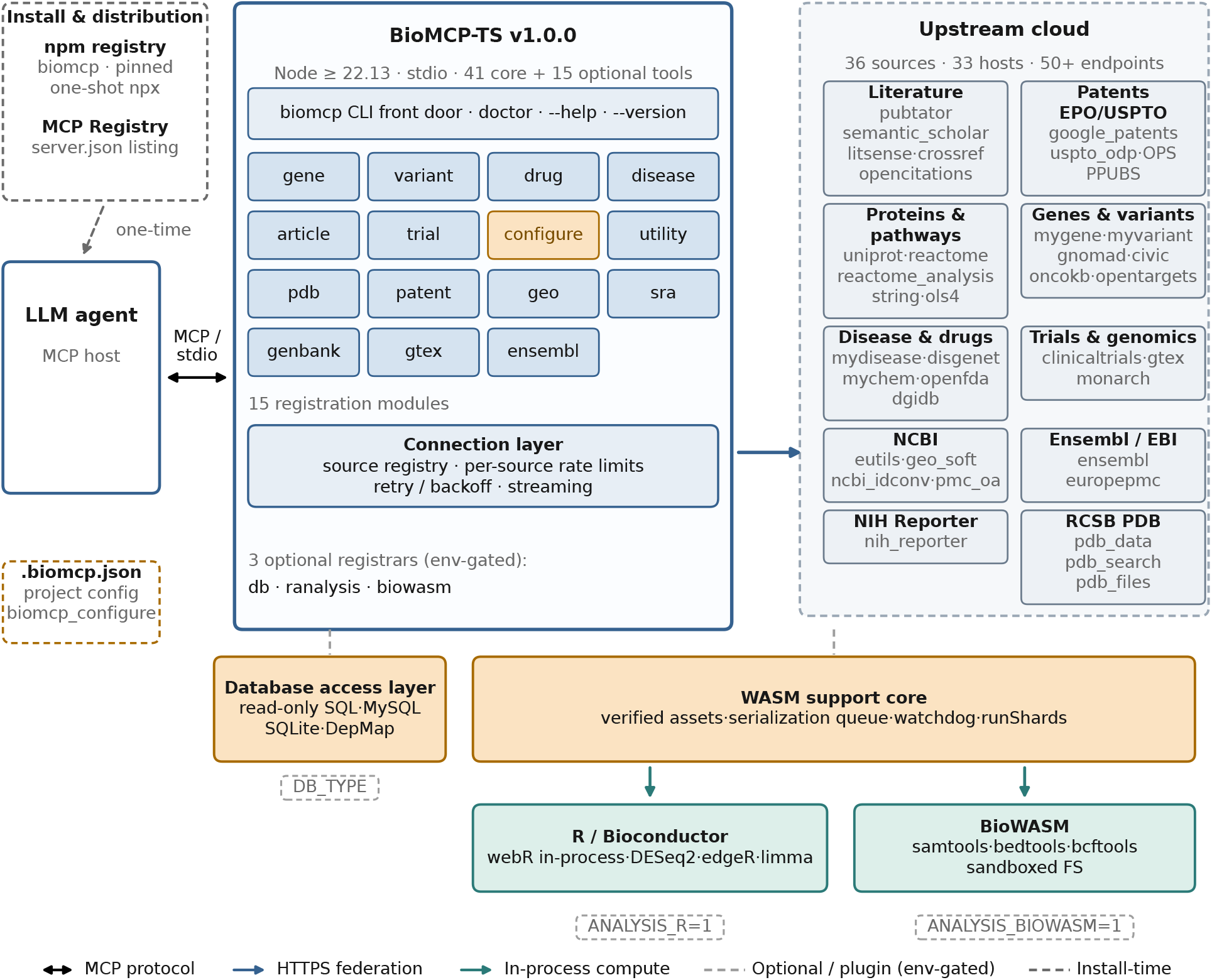
Architecture of BioMCP-TS v1.1.0; a tool is a function the agent can call. Install-time (one dashed step, top left): the pinned one-shot npm command and the MCP Registry metadata reach the host once. An MCP host then launches the server through its CLI front door (doctor, --help, --version never start the stdio server) and communicates over stdio, with 15 always-registered modules (41 core tools) reaching 33 upstream data hosts through a connection layer with rate limiting and retry. Optional plugins sit on two generic layers: a read-only SQL layer (MySQL and SQLite backends; the DepMap data pipeline is a concrete instance) and a WebAssembly layer built on the shared wasmcore support core (verified assets, serialization queue, watchdog, sharded execution), executing webR in-process for Bioconductor analysis and biowasm in worker threads for htslib operations, with artifacts chained between tools through a shared virtual filesystem. The biomcp configure meta-tool observes and writes the project configuration file.

**Figure 9.**
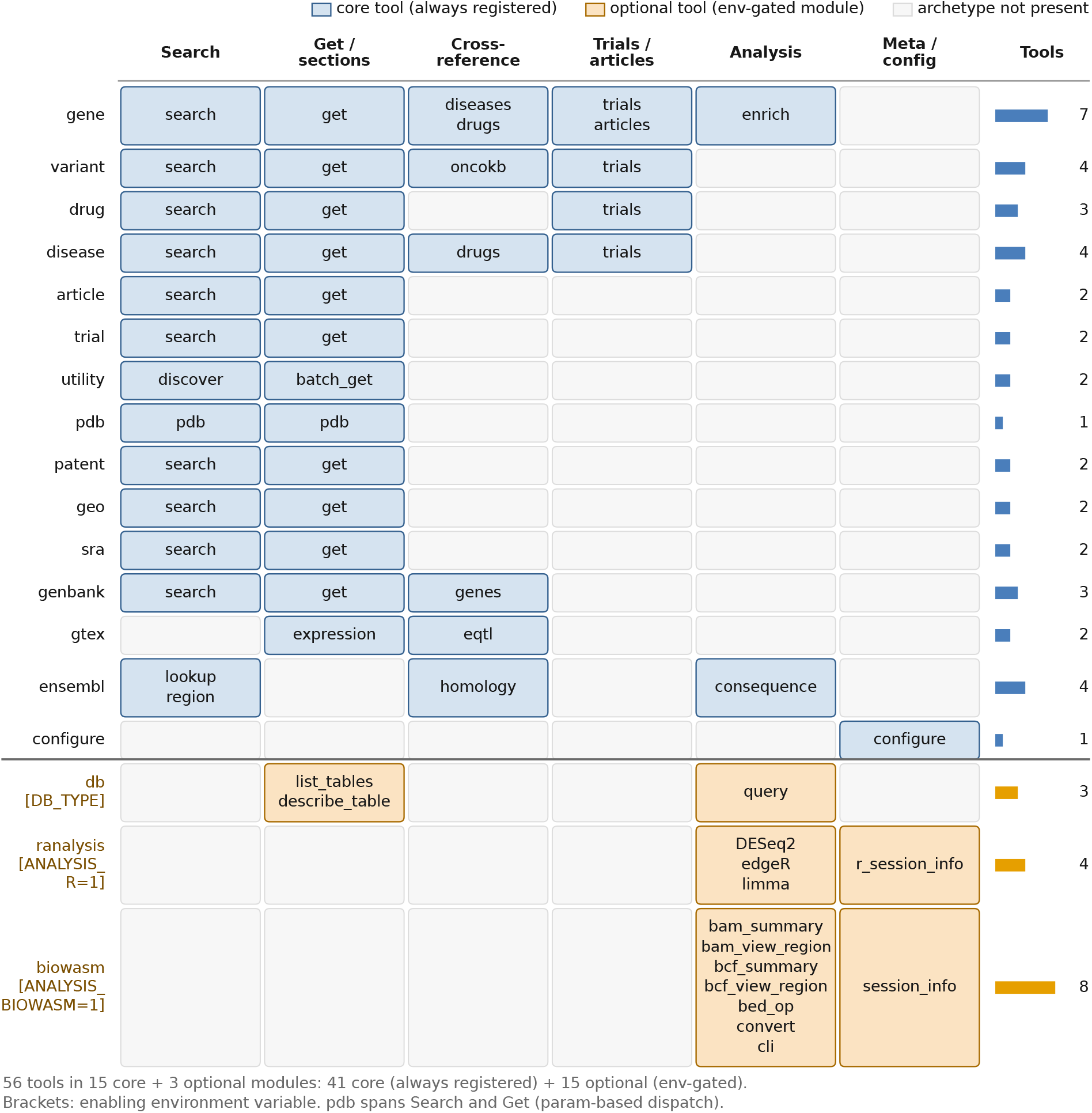
kTool inventory of BioMCP-TS v1.1.0, parsed directly from the server’s tool registry; a tool is a function the agent can call. Rows are the 15 alwaysregistered modules plus the three optional feature sets; columns group tools by archetype (search, section-based retrieval, cross-referencing, domain-specific lookup, analysis). Core tools and optional plugins are distinguished by fill; the counts strip on the right shows tools per module.

### 3.2 Federation layer

The 15 always-registered modules cover genes, variants, drugs, diseases, articles, trials, structures, patents, and functional genomics. Upstream connections run through a registry of 36 API sources with per-source rate limiting and retry; two patent backends are managed at the entity layer (keyless USPTO Public Search as the default United States backend, keyed EPO OPS for worldwide coverage). Retrieval follows a uniform contract: search tools resolve free text to canonical identifiers, and retrieval tools accept an identifier plus optional sections, fanning out with an 8 s per-section timeout; a failed source returns a structured error field for that section instead of failing the call (visible as recorded errors in Cases 2, 3, and 5). Federated article search queries five literature backends concurrently and deduplicates by PMID, PMCID, and DOI, with a fast and a full citation mode. Patent search performs co-citation mining for seminal prior art by default (the mechanism behind Case 2’s WO1995/011699 discovery). Optional API keys raise rate limits or unlock premium sources, and responses tag which source served each field, so keyless fallbacks are visible rather than silent (Case 1’s drug lane).

### 3.3 Plugin layers and the registration contract

A registration module is a function invoked once at server start that registers its tools with the SDK; the 15 core registrars always run, and each optional registrar runs only when its environment flag is set (`DB_TYPE, ANALYSIS R=1, ANALYSIS BIOWASM=1`), so inactive plugins never appear in the advertised schema (Case 4’s server advertised 44 tools).

#### Database layer

Optional SQL tools execute read-only statements through a keyword allow-list with literal-stripping scans that block writes, multi-statement input, and export clauses; MySQL and local-file SQLite backends are supported, and comma-listed SQLite files attach under derived aliases for cross-database joins. The DepMap ETL builds the Case 4 reference database from nine md5-verified release files.

#### WebAssembly layer

Both compute plugins share wasmcore, a runtime-agnostic support core: sha256-verified asset downloads, a local mirror server, a serialization queue, memory watch and watchdogs, progress throttling, and sharded bounded-concurrency execution. The webR lane runs Bioconductor (DESeq2/edgeR/limma, 63-package closure) on R 4.6.0 compiled to wasm32 inside the server process; inputs are raw integer count matrices (up to 50,000 genes by 64 samples) with sample metadata and a whitelisted design formula. The biowasm lane runs pinned samtools, bedtools, and bcftools builds in worker threads with a pool, progress notifications, cancellation, and content-addressed artifact handles (identifier, host path, sha256, preview, payloads up to 2 MB inlined); handles are accepted as inputs by later calls and chain through a per-session shared virtual filesystem, which is what lets one tool’s output file become another tool’s input (Case 5). A large-input gate requires explicit acknowledgment for full-file scans, and progress-bearing calls keep the client connection alive through long operations. In the development probe, a long scan completed against a reference client’s much shorter request timeout when progress was reported, and was dropped at that deadline when silent.

#### CLI front door

The installed `biomcp` binary is a CLI module first: `--help`, `--version`, and `biomcp doctor` never start the stdio server, and any unrecognized argv hands off to the server bundle unchanged, so every existing client configuration behaves identically. `doctor` reproduces the server’s startup (configuration resolution and feature gates) and reports a self-diagnostic: the Node engines gate, install-mode-specific advice, configuration-file health, feature gates with peer-dependency resolvability, masked environment presence, and structured `blockers` each carrying a paste-ready fix command; `doctor` exits 1 if any blocker fires. Case 0 records the practitioner-facing outcome: install, verify, first answer. The server also exits when its client closes stdin, so an abruptly killed client (SIGKILL of the parent) no longer strands a server process and its ∼1 GB webR worker holding pipes open.

#### Distribution and install pinning

The package ships a `server.json` registration in the MCP Registry format (`io.github.yeyuan98/biomcp-ts`, npm package mapping, and environment-variable documentation with secrets flagged), so agent-side tooling can resolve and install it from the Registry rather than from prose; the listing was live at submission time (the Registry is shared infrastructure and other biomedical servers are listed too; the differentiators claimed here are the self-diagnostic CLI and the install commands rendered from source). The recommended zero-install client command is generated from the release itself: the version pins (biomcp@1.0, webr@0.6, mysql2@3) are rendered from `src/version.ts` and the package’s peer-dependency ranges, with drift-guard tests enforcing that the documented commands, the Registry metadata, and the published version never disagree. A three-way name collision on “BioMCP” (the Python/Rust `biomcp` by genomoncology, and an unrelated Registry listing of the same title) makes this nameresolution-time disambiguation part of the deployment story rather than documentation politeness.

#### Configuration observability

The always-registered `biomcp configure` meta-tool observes every parameter with status and provenance and reports per-feature settable keys, so an agent can construct a valid change from an empty state. The full parameter catalog moved behind a `filter` argument (shrinking the default status payload by about 60%); writes go to an atomic 0600-mode project configuration file with secret values redacted, environment variables are strictly query-only with masked values, and the tool explains the restart and verification steps, so agents can self-serve the optional feature sets without editing environment blocks by hand.

### 3.4 Testing patterns

The verification record spans three tiers. Unit: 1,313 mocked tests in 78 suites. Integration: 181 declared live-API tests in 19 files against the real upstream services. Agent-level end-to-end: a harness drives the bundled server through a real agent CLI (opencode 1.18+) on 18 scenario suites, 14 of them usability levels L0–L3 (from first-contact orientation to multi-step error recovery, including an impossible-task honesty probe) and 4 configuration scenarios, graded by a 12-check objective vocabulary (tool-sequence, JSON-path, no-such-tool, and host-bypass probes) with written rubrics adjudicated manually where intent is subjective. The latest recorded round stands at 18 of 18 passing (round 4, recorded at v0.8.0, 2026-08-30), and agent-usability claims in this report are scoped to that single-client setup. Continuous integration gates typecheck, mocked tests, build, and a lowest-severity dependency audit. The synthetic planted-gene recovery demo (40 of 40 planted genes recovered by the webR lane) is retained as a test oracle for the compute pipeline, not as a showcase. The seven cases in this report were recorded with the same runner used by the problem set, and the pre-registered protocol (amended before recording) fixes measurement mechanics, branches, and honesty rules; all transcripts preserve errors and tool-side truncation verbatim.

## 4. Limitations and boundaries

### Scale and parity

All numbers are indicative, single-host development measurements on one workstation (Windows Subsystem for Linux 2, Ryzen 7 9700X, 16 cores, 15 GiB RAM), pinned to v1.1.0; they are not competitive benchmarks. The recording host has no native htslib installation (the deployment situation this server targets), so the cohort-VCF full-scan figure (0.8 MB s^−1^) is an absolute value and no overhead is attributed specifically to WebAssembly. Numerical parity between the webR lane and native Bioconductor installations was not assessed; Case 6 validates the compute lane end to end on real data (and cold/warm byte-identity), not bit-level agreement with native runs. The DE lane caps inputs at 50,000 genes by 64 samples; Case 4’s executed build is the full nine-file reference database, and the ETL also supports smaller subsets.

### Upstream realities

The server is read-only and inherits the rate limits, terms, and availability of its upstream data hosts; live-network timing varies substantially across runs. Optional keys change behavior (for example, DisGeNET associations replace the keyless OpenTargets fallback) in ways that response tagging makes visible but cannot eliminate. Specific environment findings were recorded verbatim in the transcripts: PDB text search misses known structures, so the identifier path is taught explicitly (Case 1); the WebAssembly runtime download carries a configurable timeout whose errors include a self-fetch and local-mirror recipe, and the server declined one full cohort-VCF scan under its large-input gate (Case 5 takes the documented slice workaround); and one known engine bug (combining an expression filter with varianttype projection in the VCF region tool) is avoided and documented rather than hidden.

### Data and deployment boundaries

DepMap staging is manual by design (CAPTCHA-gated portal; nine files md5-verified; CC BY 4.0 with portal terms for commercial use). Case 5’s fixtures are a low-coverage NA12878 alignment and a 2,504-sample cohort VCF on different chromosomes, so coverage is sample-side and cohort allele frequencies serve as population annotation, with no genotype-concordance claims. Because an agent may consume untrusted content, the threat surface is bounded but not zero: the configuration meta-tool writes only its own project file (atomic, mode 0600, secrets redacted), and the SQL layer attaches only explicitly configured paths under a read-only allow-list; agents handling untrusted documents should still run with least privilege.

### Scope and sustainability

The landscape positioning (Table 1) reflects a scan conducted in September 2026 of MCP server distributions on npm and GitHub, the MCP Registry API, preprint indexes (arXiv), and the cited publications, restricted to packages whose documented scope is bioinformatics; competitor attributes were not benchmarked, and the negative compute claim retains its “to our knowledge” scope. Singlecell analytics and genome-region triage beyond Case 5’s pattern are adjacent applications not covered by this release. The MCP ecosystem is young and its conventions are still moving; the protocol surfaces used here (tools, schemas, stdio transport, progress) are stable, but ecosystem practices for sandboxing and provenance will keep evolving. BioMCP-TS is derived in part from the open-source BioMCP project (MIT license) (Genomoncology, 2025); the TypeScript implementation, tool registry, plugin layers, and evaluation described here are original to this work. Single-maintainer sustainability is a disclosed risk; the three-tier verification record and the agent-level harness exist precisely to lower the cost of future maintenance.

## Availability

Project and source (Apache-2.0):https://github.com/yeyuan98/biomcp-ts. Install (minimal, version-pinned): npx -y -p biomcp@1.1.0 biomcp; the all-features one-shot adds the analysis peers (-p webr@0.6-p mysql2@3); verify with npx -y -p biomcp@1.1.0 biomcp doctor (Node ≥ 22.13; zero configuration). The server is listed in the MCP Registry (`io.github.yeyuan98/biomcp-ts`) for agent-side discovery.

The exact release evaluated here is the npm package biomcp@1.1.0 (tarball SHA-1 `fbb086fa`). The practical-case problem set (verbatim prompts, tool-call transcripts, runner scripts, expected-answer snapshots, compute-tracking raw outputs, and fixture pointers with checksums) is published as one Zenodo dataset: https://doi.org/10.5281/zenodo.22279669 (Yuan, 2026a) (this evaluation’s version; the concept DOI 10.5281/zenodo.22178967 always resolves to the latest, and the dataset retains the v0.8.0 and v1.0.0 trace generations alongside v1.1.0 for cross-reference). Sequencing fixtures are published under CC0 (Zenodo DOI 10.5281/zenodo.22156404). A versioned software archive DOI for v1.1.0 is https://doi.org/10.5281/zenodo.22279640 (Yuan, 2026b) (concept DOI 10.5281/zenodo.22178957). DepMap data are used under CC BY 4.0 (Broad Institute DepMap Portal, 2026).

## Supplementary inventory

**S1** Problem set and runner scripts, including verbatim prompts, full tool-call transcripts, answers, and preserved errors for Cases 0–6 (Zenodo dataset).

**S2** Pre-registered benchmark protocol and its amendments A2/A3, with raw per-run outputs.

**S3** Figure source data and QA records for all figures.

**S4** Claim-to-evidence mapping linking every quantitative statement in the text to repository evidence or recorded artifacts.

## Acknowledgements

The author thanks the biowasm and webR projects for WebAssembly builds of the samtools, bedtools, bcftools, and R toolchains, the upstream BioMCP project for the seed of this implementation, the Bioconductor and DepMap teams for open data and software, and the maintainers of the many public APIs this server federates.

## Author contributions

Y.Y.: Conceptualization, Methodology, Software, Validation, Investigation, Data curation, Visualization, Writing (original draft), Writing (review and editing).

